# Transcription factor activity rhythms and tissue-specific chromatin interactions explain circadian gene expression across organs

**DOI:** 10.1101/207787

**Authors:** Jake Yeung, Jérôme Mermet, Céline Jouffe, Julien Marquis, Aline Charpagne, Frédéric Gachon, Félix Naef

## Abstract

Temporal control of physiology requires the interplay between gene networks involved in daily timekeeping and tissue function across different organs. How the circadian clock interweaves with tissue-specific transcriptional programs is poorly understood. Here we dissected temporal and tissue-specific regulation at multiple gene regulatory layers by examining mouse tissues with an intact or disrupted clock over time. Integrated analysis uncovered two distinct regulatory modes underlying tissue-specific rhythms: tissue-specific oscillations in transcription factor (TF) activity, which were linked to feeding-fasting cycles in liver and sodium homeostasis in kidney; and co-localized binding of clock and tissue-specific transcription factors at distal enhancers. Chromosome conformation capture (4C-Seq) in liver and kidney identified liver-specific chromatin loops that recruited clock-bound enhancers to promoters to regulate liver-specific transcriptional rhythms. Furthermore, this looping was remarkably promoter-specific on the scale of less than ten kilobases. Enhancers can contact a rhythmic promoter while looping out nearby nonrhythmic alternative promoters, confining rhythmic enhancer activity to specific promoters. These findings suggest that chromatin folding enables the clock to regulate rhythmic transcription of specific promoters to output temporal transcriptional programs tailored to different tissues.

## Introduction

A mammalian internal timing system, known as the circadian clock, orchestrates temporal physiology in organs to anticipate daily environmental cycles (Dibner & Schibler 2015). Individual cells within organs contain a molecular oscillator that, together with rhythmic systemic signals such as hormones, temperature, and feeding behavior, collectively drive diurnal oscillations in gene expression and physiology (Lamia et al. 2008; Reinke et al. 2008; Vollmers et al. 2012; Cho et al. 2012). Remarkably, the circadian clock impinges on many gene regulatory layers, from transcriptional and posttranscriptional processes, translation efficiency, to translational and posttranslational processes (Mermet et al. 2016).

Transcriptome analysis of large collections of mammalian cell types and tissues have highlighted the breadth of tissue-specific transcriptional regulation (Yue et al. 2014; Merkin et al. 2012). However, many physiological processes are dynamic at the timescale of hours and often under circadian control, such as hormone secretion, drug and xenobiotic metabolism, and glucose homeostasis (Takahashi et al. 2008). Therefore, unlocking the temporal dimension to tissue-specific gene regulation is needed for an integrated view of physiological control.

Chronobiology studies have shown that different tissues utilize the circadian clock to drive tissue-specific rhythmic gene expression (Storch et al. 2002; Zhang et al. 2014; Korenčič et al. 2014), presumably to schedule physiological functions to optimal times of day. Indeed, genetic ablation of the circadian clock in different tissues can lead to divergent pathologies, such as diabetes in pancreas-specific *Bmal1* knockout (KO) and fasting hypoglycemia in liver-specific *Bmal1* KO, suggesting that the clock interweaves with tissue-specific transcriptional programs (Bass & Lazar 2016). But how diurnal and tissue-dependent regulatory landscapes interact to generate tissue-specific rhythms is poorly understood.

## Results

### Contributions of tissue, daily time, and circadian clock to global variance in mRNA expression

To estimate the respective contributions of tissues, daily time, and circadian clock to global variance in gene expression, we analyzed available temporal transcriptomes across 11 tissues in WT mice (Zhang et al. 2014), and generated temporal RNA-Seq data of liver and kidney from *Bmal1* KO mice and WT littermates (Supplemental Table S1 & Supplemental Table S2, Methods). The Zhang *et al.* dataset was obtained under dark-dark (DD), ad libitum feeding, sampled every 2 hours. The liver and kidney *Bmal1* KO and WT datasets were obtained under light-dark, night-restricted feeding (LD) conditions, sampled every 4 hours.

To avoid mixing different experimental designs (e.g. temporal resolution and number of repeats, Deckard et al. 2013; Li et al. 2015), we analyzed these datasets separately. We performed principal component analysis (PCA) on the entire set of conditions (11 tissues times 24 time points) to obtain a first unbiased overview into the contributions of tissue and time-specific variance in the data. This showed that most of the variance concerned differences in expression between tissues (Figure 1A & Supplemental Figures S1A-D). Temporal variance, in particular 24h periodicity, was present among a group of principle components carrying lower amounts of variance (Figure 1A & Supplemental Figures S1E-G). Focusing on genome-wide temporal variation within each tissue, we found that 24-hour rhythms constituted the largest contribution of temporal variance, followed by 12-hour rhythms, which were close to background levels for many tissues (Figure 1B) (Hughes et al. 2009). We thus focused the rest of our analysis on 24h rhythms.

**Figure 1.**
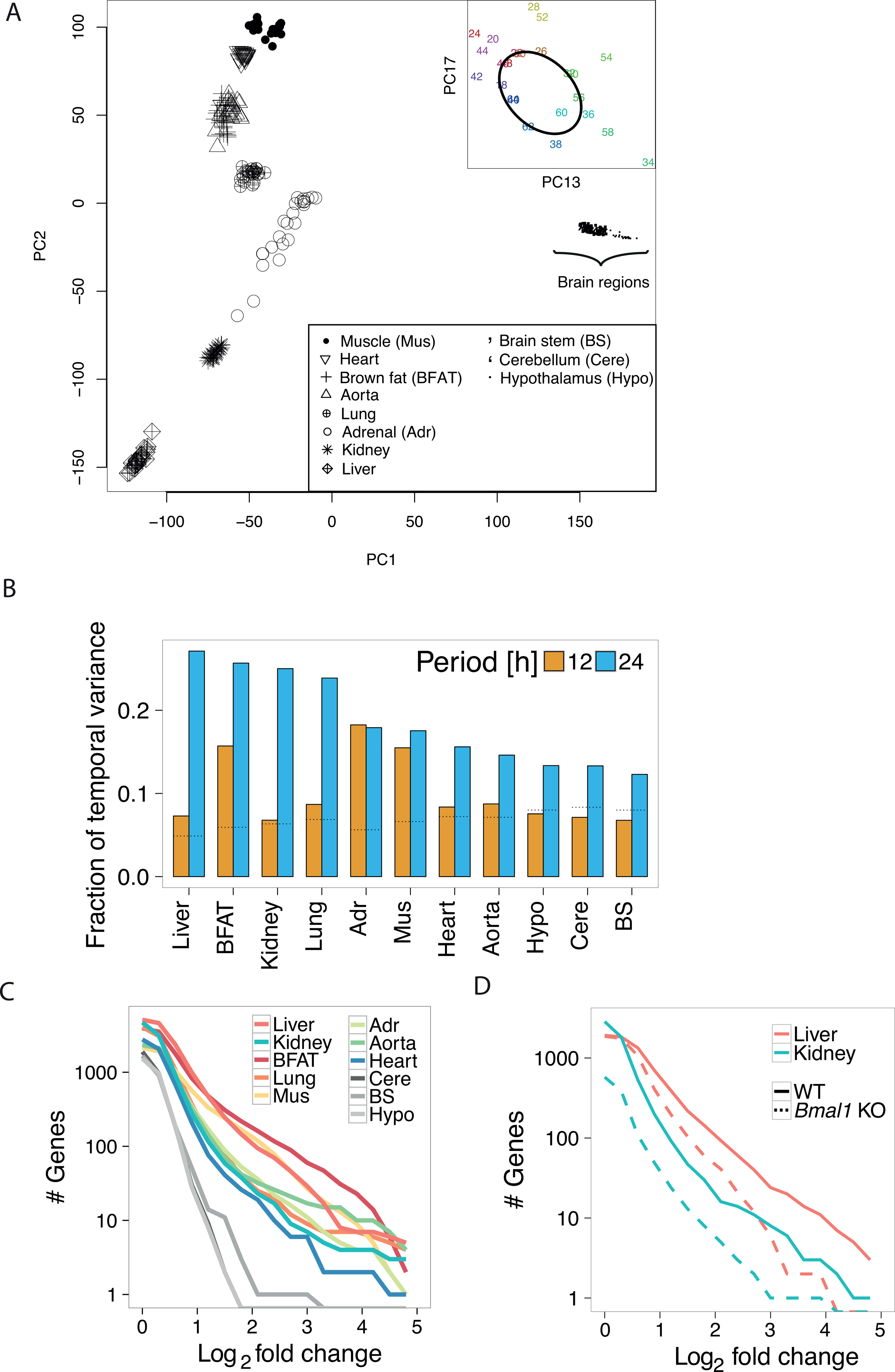
Contribution of tissue, daily time, and circadian clock to global variance in mRNA expression. (A) Principal component (PC) analysis of two days temporal transcriptomes across 11 WT tissues. PC1 and PC2 show clustering of samples by tissues; each point represents a tissue sample (see legend) at a specific time point (not labeled). Inset: Loadings for PC13 and PC17 for the liver samples labeled with circadian time (CT), showing temporal variation along an elliptic path. Colors: CT; CT0 corresponds to subjective dawn; CT12 corresponds to subjective dusk. (B) Fractions of temporal variance in each tissue explained by 24- and 12-hour periods, obtained by applying spectral analysis genome-wide for each tissue. Dotted horizontal lines represent the expected background level, assuming white noise. (C,D) Cumulative number of rhythmic genes (p<0.01, harmonic regression) with log2 fold change larger than the value on the x-axis. (C) Analysis on 11 WT tissues. (D)Analysis on 4 conditions: *Bmal1* KO mice and WT littermates in liver and kidney.

We analyzed the peak-to-trough amplitudes (hereafter also referred to as fold change) of 24h rhythmic transcripts. This showed that metabolic tissues, notably liver, brown fat, and skeletal muscle stand out as exhibiting far more (on the order of 100 transcripts) intermediate to high amplitude (between 2 and 10 fold) transcript rhythms. Brain tissues show virtually no rhythmic transcripts above 4 fold (Figure 1C). In liver and kidney of *Bmal1* KO mice, the number of rhythmic mRNAs was reduced by 3 fold compared to WT littermates. This effect increased for larger amplitudes. Only few transcripts in tissues of *Bmal1* KO oscillated by more than 10 fold (Figure 1D). Thus, a functional circadian clock is required for high amplitude transcript rhythms across diverse tissues, while systemic signals regulate lower amplitude rhythms that persist in clock-deficient liver (Hughes et al. 2012; Atger et al. 2015; Sobel et al. 2017) and kidney (Nikolaeva et al. 2012).

### Combinatorics of rhythmic transcript expression across tissues and genotypes

We reasoned that identifying sets of genes with shared rhythms across subsets of tissues would allow finding underlying regulatory mechanisms. We therefore developed a model selection (MS) algorithm extending harmonic regression (Fisher 1929) to classify genes into modules sharing rhythmic mRNA profiles across subsets of tissues (Figure 2A, Methods). Phase-amplitude relationships (phase is defined as the time of the peak, and amplitude as the log2 fold change) between genes and tissues are summarized using complex-valued singular value decomposition (SVD) (Figure 2B, Methods). We applied MS to the 11 tissues, which identified gene modules involving rhythmic mRNA accumulation in nearly all tissues (tissue-wide) (Figure 2C), in single tissues (tissue-specific), or in several 5 tissues (tissue-restricted) (examples shown in Figure 2D & Supplemental Figure S2A & Supplemental Table S3).

**Figure 2.**
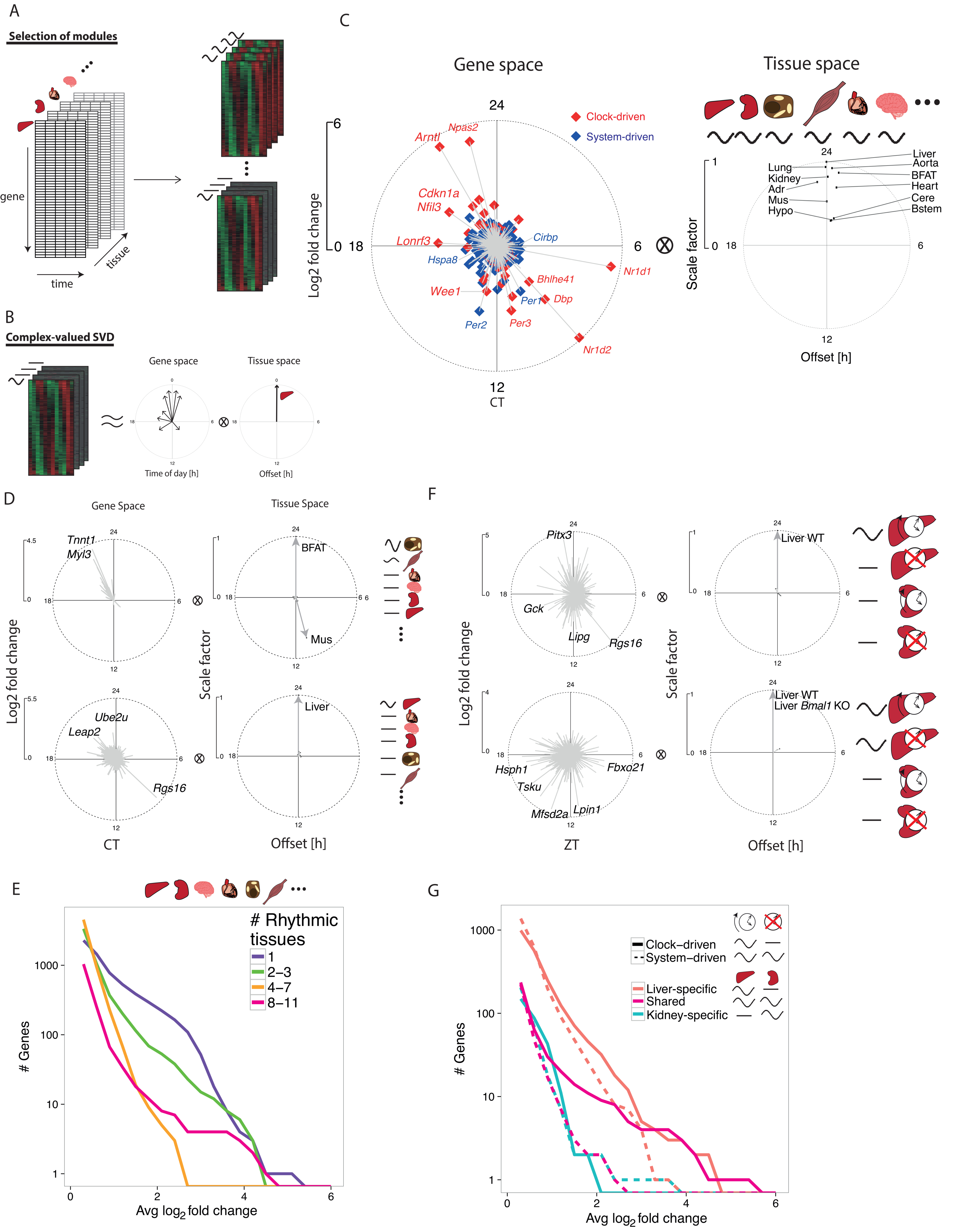
Combinatorics of rhythmic transcript expression across tissues and genotypes. (A) Schema for the model selection (MS) algorithm to identify rhythmic gene expression modules across tissues. Temporal transcriptomes of different tissues represented as a 3-dimensional array (left). Gene modules are probabilistically assigned amongst different combinations of 24-hour rhythms across tissues (e.g. tissue-specific or tissue-wide rhythms schematically shown on right). (B) Gene modules are summarized by the first component of complex-valued singular value decomposition (SVD) to highlight phase (peak time shown as the clockwise angle) and amplitude (log2 fold change shown as the radial distance) relationships between genes (gene space) and between tissues (tissue space). SVD representation is scaled such that the genes show log2 fold changes, while tissue vectors are scaled such that the highest amplitude tissue has length of 1 and a phase offset of 0 hours. (C-E) MS applied to 11 WT tissues. (F,G) MS applied to *Bmal1* KO and WT littermates in liver and kidney. (C) SVD representation of tissue-wide mRNA rhythms from the 11 tissues. Genes are labeled as system-driven (blue) or clock-driven (red) according to the comparison of the corresponding temporal profiles in *Bmal1* KO and WT littermates (Methods). (D) Examples of anti-phasic rhythms (brown fat and muscle, n=20, first SVD component explains 81% of variance), and tissue-specific rhythms (liver, n=846, first SVD component explains 59% of variance). Representative genes with large amplitudes are labeled. (E) Number of transcripts showing rhythms (p-value < 0.01, harmonic regression) in different numbers of tissues, in function of increasing peak to trough amplitudes on the x-axis. X-axis: average log2 fold change calculated from the identified rhythmic tissues. (F) SVD representation of clock- (top, n=991, 83% of variance) and system-driven (bottom, n=1395, 84% of variance) liver-specific rhythms. Number of transcripts showing clock- (solid) or system-driven (dotted) rhythms (p-value < 0.01, harmonic regression) in liver (red), kidney (blue), or both (magenta).

The tissue-wide module contained a set of both clock- and system-driven rhythmic mRNAs, as determined by comparing *Bmal1* KO data in liver and kidney (Figure 2C, left). Moreover, these transcripts oscillated in synchrony across all tissues and peaked at fixed times of day, albeit their amplitudes varied between tissues, with brain regions showing the smallest amplitudes (Figure 2C, right). The clock drove synchronized oscillations at high amplitudes, notably clock genes (e.g. *Arntl, Npas2, Nr1d1,2*; note that *Arntl* and *Nr1d1,2* are also named *Bmal1* and *Rev-erba,b* respectively), clock output genes (e.g. *Dbp, Nfil3*), and cell cycle regulators (*Cdknla* and *Weel*) (Gréchez-Cassiau et al. 2008; Matsuo et al. 2003). Interestingly, clock genes *Per1,2* continued to oscillate in *Bmal1* KO in multiple tissues, extending previous studies in liver (Kornmann et al. 2007). Other clock-independent oscillations included mRNAs of heat- and cold-induced genes, such as *Hspa8* and *Cirbp* (Morf et al. 2012; Gotic et al. 2016), that peaked 12 hours apart near CT18 and CT6 (CT: circadian time; CT0 corresponds to subjective dawn and start of the resting phase; CT12 corresponds to subjective dusk and start of the activity phase), concomitantly with highs and lows in body temperature rhythms (Refinetti & Menaker 1992).

Tissue-restricted modules contained rhythmic transcripts that peaked in synchrony, such as in liver and kidney, or with fixed offsets, such as the nearly 12 hours shifted rhythms in brown fat and skeletal muscle (Supplemental Figure S3A). Overall, transcripts with large amplitudes (FC>8) oscillated in either a few tissues (3 or less) or tissue-wide (8 or more) (Figure 2E).

To distinguish clock- and system-driven mRNA rhythms, we applied the MS algorithm to the liver and kidney transcriptomes in WT and *Bmal1* KO mice (Figure 2F & Supplemental Figure S3B & Supplemental Table S4). This separation identified clock- and system-driven modules that oscillated in liver but were flat in kidney (Figure 2F), as exemplified by mRNAs of *Lipg* and *Lpin1* (Supplemental Figure S2B). Indeed, both transcripts oscillated in WT liver with robust amplitudes, peaking near ZT11, but were flat in kidney (ZT: Zeitgeber time; ZT0 corresponds to onset of lights-on; ZT12 corresponds to onset of lights-off). However, in *Bmal1* KO, *Lpin1* continued to oscillate, while *Lipg* was flat.

Summarizing, we found that shared clock-driven mRNA rhythms, which contained core clock and clock-controlled genes, oscillated with significantly larger amplitudes than system-driven genes (Figure 2G, magenta solid versus dotted). Similarly, clock-driven liver-specific mRNA rhythms also oscillated at higher amplitudes compared with system-driven mRNA rhythms (Figure 2G, red solid versus dotted). On the other hand, kidney-specific clock- and system-driven transcripts oscillated with comparable amplitudes (Figure 2G, blue solid versus dotted), and were less numerous overall, which could reflect the distinct cell types constituting the kidney (Lee et al. 2015). The uncovered diversity of clock- and system-driven mRNA rhythms involving distinct combinations of tissues hints at complex transcriptional or post-transcriptional regulation. Below, we examine transcription regulators responsible for tissue-specific mRNA rhythms.

### Oscillatory TF activity in one tissue but not others can drive tissue-specific mRNA rhythms

We focused on WT and *Bmal1* KO liver and kidney to identify rhythmic TF activities underlying clock- and system-driven tissue-specific mRNA rhythms. We first analyzed liver-rhythmic genes driven by systemic signals (n=1395, MS; Figure 3A), which were associated with feeding and fasting rhythms (GO analysis around the clock, Method). Indeed, ribosome biogenesis was upregulated most strongly during the first six hours of the feeding phase (from ZT12 to ZT18) (Jouffe et al. 2013; Chauvin et al. 2014), while insulin signaling was downregulated during first six hours of the fasting phase (from ZT0 to ZT6) (Ravnskjaer et al. 2013), consistent with daily responses to nutrient fluctuations in liver (Sinturel et al. 2017).

**Figure 3.**
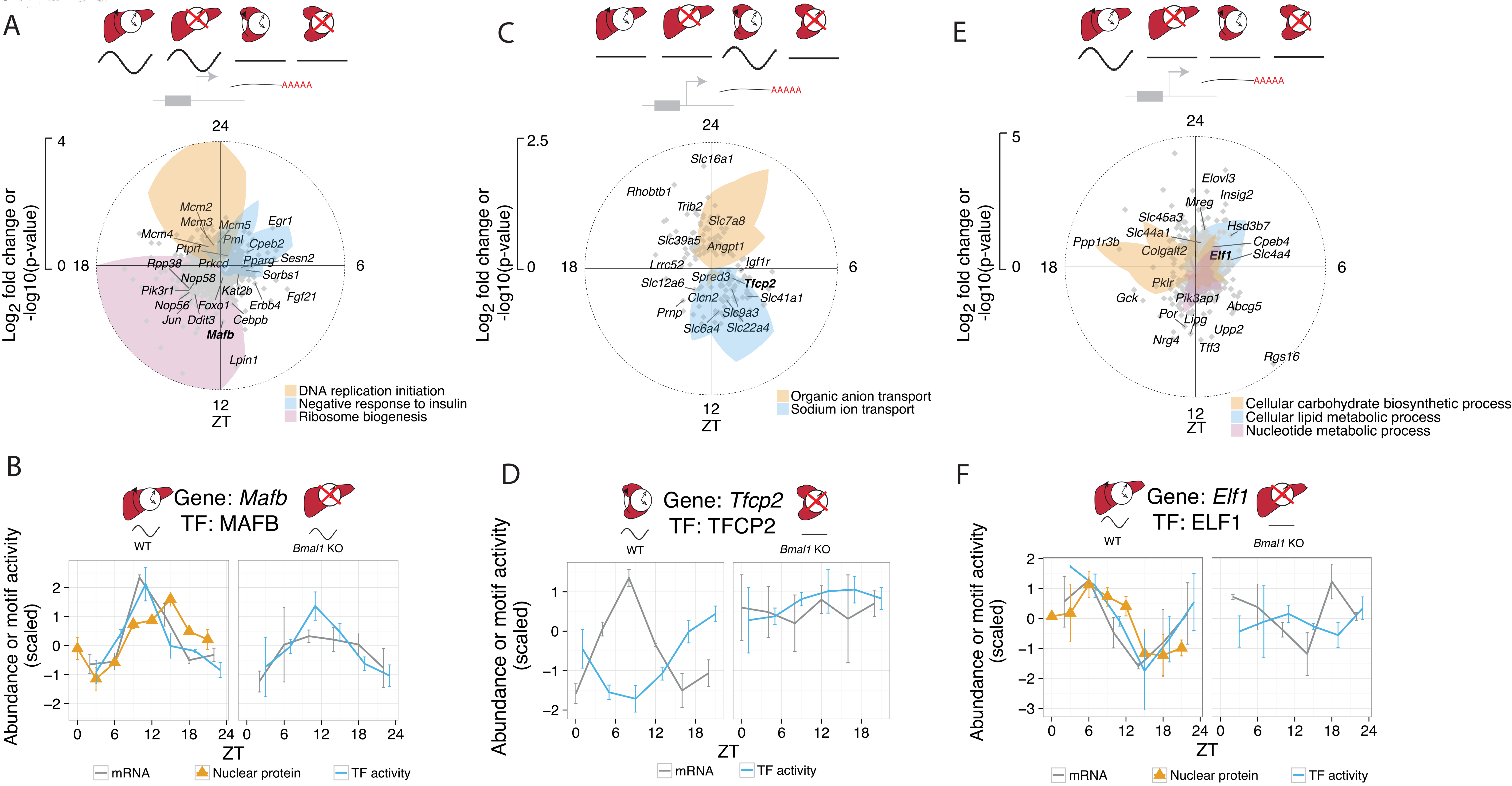
Oscillatory TF activity in one tissue but not others can drive tissue-specific rhythms. (A) Module describing system-driven liver-specific rhythms (n=1395, first SVD component explains 84% of variance). Radial coordinate of the colored polygons represent enrichment of the indicated GO terms at each time point, obtained by comparing the genes falling in a sliding window of +/− 3 hours to the background set of all 1395 genes assigned to module (p-value computed from Fisher’s exact test). (B) MAFB is a candidate TF for the module in A. Predicted MAFB activity (solid line), nuclear protein abundance (triangles), and mRNA accumulation (dotted) oscillate in WT and *Bmal1*&KO, with peak mRNA preceding peak nuclear protein and TF activity. Error bars in nuclear protein, mRNA, and TF activity show SEM (n=2). (C) Clock-driven kidney-specific module (n=156, first SVD component explains 80% of variance). Colored polygons as in (A). (D) TFCP2 is a candidate TF for the module in C. The temporal profile of predicted TFCP2 activity (solid line) is anti-phasic with *Tfcp2* mRNA accumulation (dotted) in WT, and both are flat in *Bmall* KO. Error bars in mRNA and TF activity show SEM (n=2). (E) Clock-driven liver-specific module (n=991, first SVD explains 83% of variance). (F) ELF is a candidate TF for the module in E. The temporal profile of predicted ELF activity (solid line) in WT matches that of nuclear protein abundance in liver (triangles), and both are delayed compared to *Elfl* mRNA accumulation (dotted). In *Bmal1* KO, ELF activity and *Elfl* mRNA are nonrhythmic. Error bars in nuclear protein, mRNA, and TF activity show SEM (n=2).

To infer rhythmic TF activities that may underlie these mRNA rhythms, we applied a penalized regression model (MARA) (Balwierz et al. 2014) that integrates TF binding site predictions near promoters with mRNA accumulation. TF analysis of this module notably identified TFs related to insulin biosynthesis and gluconeogenesis, such as MAFB (Matsuoka et al. 2003) and EGR1 (Matsuoka et al. 2003; Shen et al. 2015), whose activities peaked at ZT11 and ZT3, respectively (Figure 3B & Supplemental Figure S4A). Integrating temporal activities of candidate TFs with RNA-Seq and our previously described temporal nuclear protein dataset (Wang et al. 2017), we found that rhythmic activity of MAFB and EGR1 was supported by rhythmic mRNA abundance followed by rhythmic nuclear protein abundance (Figure 3B, Supplemental Figure S4B), likely reflecting the delayed protein abundance after mRNA accumulation (Mermet et al. 2016).

Next, we analyzed clock-driven transcripts oscillating specifically in the kidney (n=156, MS; Figure 3C), among which sodium ion and organic anion transporters peaked near ZT12 and ZT0, respectively. The upregulation of sodium ion transporters in kidney during the behaviorally active phase may underlie clock-dependent increase of sodium excretion (Nikolaeva et al. 2012). Similarly, the upregulation of organic anion transporters during the resting phase may explain increased transport activity for precursors of gluconeogenesis, such as pyruvate and lactate, during fasting (Ekberg et al. 1999; Stumvoll et al. 1998). mRNAs that peaked during the resting phase may be regulated by TFCP2, as predicted by TF analysis (Figure 3D & Supplemental Figure S4C). In addition, the predicted TFCP2 activity was anti-phasic with *Tfcp2* mRNA abundance, suggestive of a repressive activity, consistent with the ability of TFCP2 to recruit histone deacetylase HDAC1 (Kim et al. 2016).

Finally, liver-specific clock-driven rhythmic transcripts (n=991, MS) were comprised of genes associated with glucose metabolism (enriched at ZT18), such as *Gck* and *Ppp1r3b* (Kelsall et al. 2009; Oosterveer & Schoonjans 2014), as well as lipid, cholesterol, and bile acid metabolism genes (enriched at ZT2), such as *Elovl3, Insig2, Hsd3b7*, and *Cyp8b1* (Guillou et al. 2010; Le Martelot et al. 2009; Sayin et al. 2013; Shea et al. 2007) (Figure 3E). Predicted activity of ELF oscillated and peaked near ZT3 in WT liver but was flat in *Bmal1* KO (Fang et al. 2014) (Figure 3F & Supplemental Figure S4D). Interestingly, mRNA abundance of *Elf1* as well as its nuclear protein abundance also oscillated in WT, supporting *Elf1* as a potential regulator of oscillating transcriptions peaking near midday. Thus, the MS algorithm separated genes into physiologically relevant modules, allowing reliable prediction of rhythmically active TFs regulating temporal physiology of respective tissues.

### Co-localized binding of clock and liver-specific TFs drives liver-specific mRNA rhythms

To further dissect liver-specific clock-driven rhythms, we reasoned that accessible chromatin regions specific to the liver could harbor regulatory sites for clock TFs, which could then regulate mRNA rhythms liver-specifically. Comparing DNase I hypersensitive sites (DHSs) in liver and kidney (DNase-Seq data from ENCODE) (Yue et al. 2014), we found that liver-specific clock-driven genes were enriched with liver-specific DHSs (within 40 kb from promoters), compared to system-driven as well as nonrhythmic genes (Figure 4A). Using TF binding site predictions underlying these liver-specific DHSs, we applied MARA to predict rhythmic TF activities that explain gene expression of this module (Supplemental Figure S5A). In WT liver, the predicted activity of RORE oscillated with robust amplitudes and peaked near ZT21. RORE activity became high and flat in *Bmal1* KO liver, consistent with loss of REV-ERB expression and consequently derepression of REV-ERB target genes (Bugge et al. 2012) (Figure 4B, top). Activity of E-box in WT liver peaked at ZT7, consistent with BMAL1:CLOCK activity (Rey et al. 2011), albeit with weaker amplitudes compared to RORE activity, likely reflecting fewer E-box target genes compared to RORE in this module. In *Bmal1* KO mice; E-box activity was low and flat in liver, as expected.

**Figure 4.**
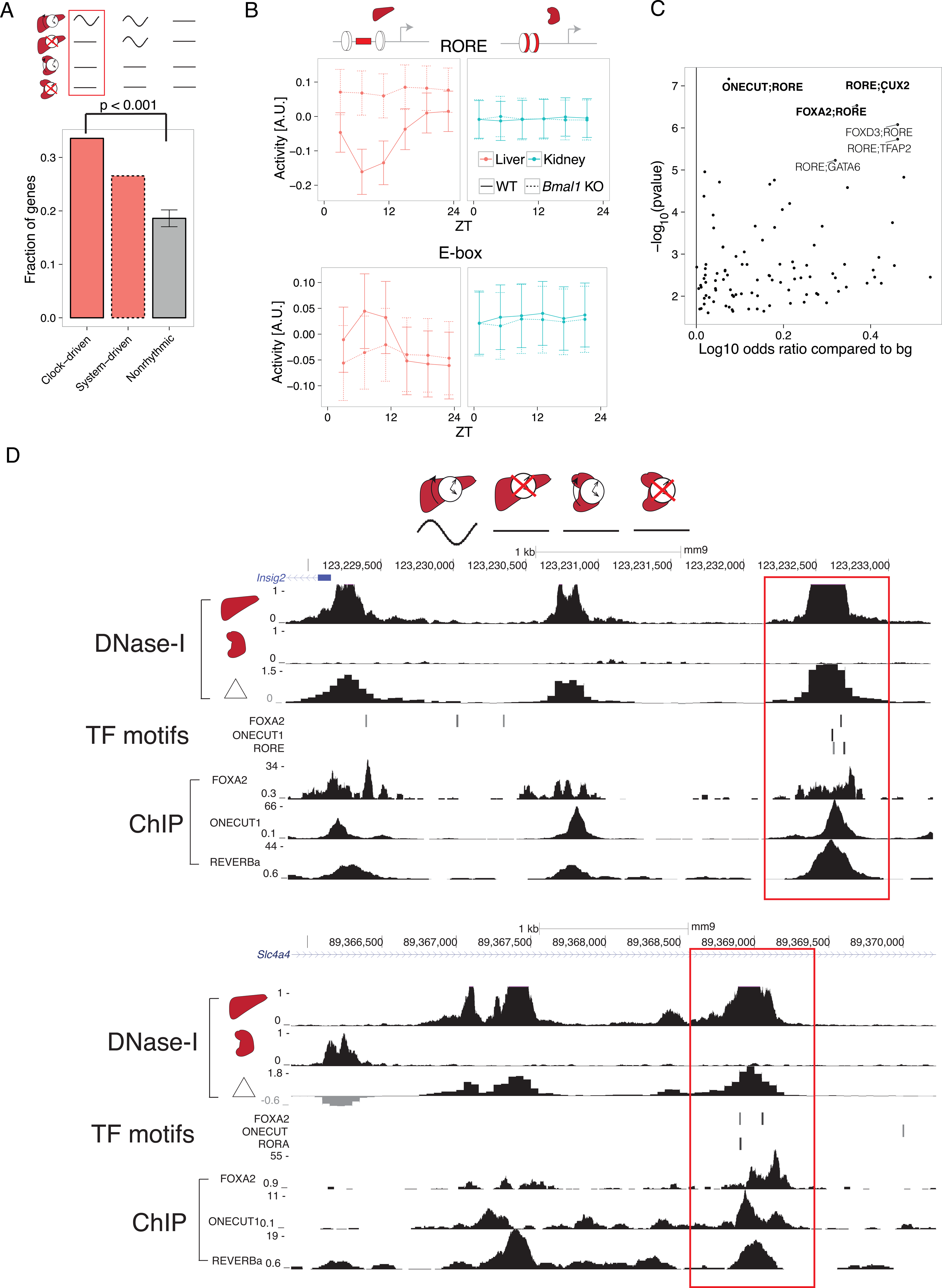
Co-localized binding of clock and liver-specific TFs underlies liver-specific mRNA rhythms. (A) The fraction of genes containing liver-specific DNase-I hypersensitive sites (DHSs) in the clock-driven liver-specific module is higher compared with both nonrhythmic and system-driven liver-specific modules. Error bars and p-values calculated from 10000 bootstrap iterations. (B) Predicted temporal activities of RORE (top) and E-box (bottom) TF motifs located within liver-specific DHSs. Error bars show standard deviation of the estimated activities. (C) Co-occurrence of RORE with all other TFs in the SwissRegulon database (189 TF motifs). Positive log10 odds ratios (ORs) represent pairs of motifs enriched in the clock-driven liver-specific module compared to the flat module. P-values for the motif pairs were calculated from chi-square tests applied to 3-way contingency tables (Myšičková et al. 2012). Selected pairs are in bold. (D) DNase-I hypersensitivity in liver, kidney, and the corresponding differential signal (in log2 fold change) near two representative genes (top: *Insig2*; bottom: *Slc4a4*). RORE, ONECUT1, and FOXA TF binding motifs (posterior probability > 0.5, MotEvo) co-occur at liver specific DHSs (red boxes). Predicted TF binding sites correspond to experimentally observed TF binding in publicly available ChIP-exo datasets for REV-ERBa, ONECUT1, and FOXA2 (bottom).

We hypothesized that cooperativity of liver-specific and clock TFs at liver-specific DHSs can regulate liver-specific mRNA rhythms. Pairwise analysis of TF binding sites at liver-specific DHSs found enrichment of co-occurrence between RORE and liver-specific TF motifs, FOXA2, ONECUT, and CUX2 (Figure 4C). Enrichment of both CUX2 and ONECUT (also named HNF6) is consistent with ONECUT1 binding to both ONECUT and CUX2 motifs (Conforto et al. 2015). mRNAs of genes with co-occurrence of RORE and liver-specific TF motifs peaked near ZT0-ZT2, consistent with peak RORE activity (near ZT21) preceding peak mRNA abundance of REV-ERB targets (Supplemental Figure S5B). Analysis of ChIP-exo datasets targeting FOXA2, ONECUT1, and REV-ERBa in liver (Iwafuchi-Doi et al. 2016; Wang et al. 2014; Zhang et al. 2015) confirmed co-localized TF binding at liver-specific DHSs distal from clock-driven liver mRNAs such as *Insig2* and *Slc4a4* (Figure 4D). Thus, colocalized binding of liver-specific and clock TFs at distal liver-specific DHSs may regulate liver-specific mRNA rhythms.

### Liver-specific chromatin loops regulate liver-specific mRNA rhythms

To test whether distally located liver-specific DHSs can contact promoters of clock-driven liver-rhythmic genes, we selected the promoters of *Mreg, Pik3ap1*, and *Slc44a1* as baits for 4C-Seq experiments in liver and kidney harvested at the time of peak mRNA accumulation for the selected genes (Methods, Figure 5A & Supplemental Figure S6A & Supplemental Figure S7A). Upstream of *Mreg*, the 4C-Seq signal, which measures frequency of promoter-enhancer contacts (van de Werken et al. 2012), decayed rapidly to background level in both liver and kidney (Figure 5B top). Downstream of *Mreg*, however, the 4C-Seq signal showed a tissue-dependent pattern, decaying slowly in the liver but more rapidly in the kidney. This difference in decay suggests increased frequency of promoter-enhancer contacts in the liver compared to the kidney. Indeed, differential analysis identified liver-specific chromatin contacts 40 kb downstream of the promoter (Figure 5B bottom). Overlaying the contact data with DNase-Seq, we found that liver-specific chromatin contacts downstream of *Mreg* connected liver-specific DHSs with the *Mreg* promoter (Figure 5C). Furthermore, ChIP-exo showed co-localization of REV-ERBa and FOXA2 binding at liver-specific DHSs contacting the promoters (Figure 5C). By contrast, accessible regions upstream of the *Mreg* promoter did not show liver-specific chromatin contacts. The 4C-Seq data thus suggest that liver-specific chromatin loops can recruit clock-bound distal elements to promoters to regulate liver-specific transcriptional rhythms. Other liver-specific rhythmic transcripts, *Pik3ap1 and Slc44a1*, also displayed liver-specific chromatin loops between promoter and liver-specific open chromatin regions (Supplemental Figure S6 & Supplemental Figure S7), corroborating that such tissue-specific looping drives tissue-specific mRNA rhythms.

**Figure 5.**
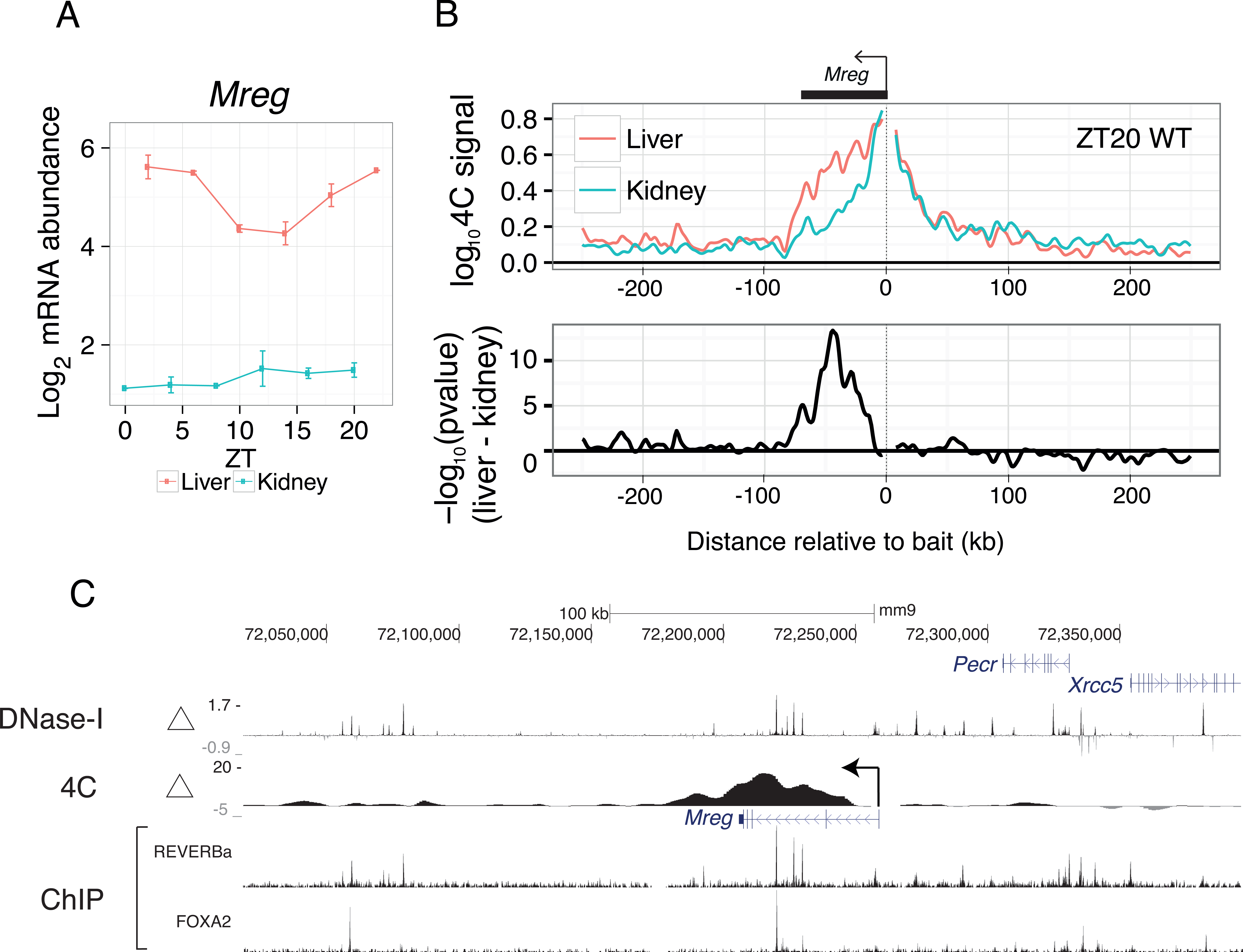
Liver-specific chromatin loops regulate liver-specific mRNA rhythms. (A) Temporal mRNA profile for *Mreg*, a clock-driven liver-rhythmic gene. Error bars are SEM (n=2). (B) 4C-Seq profiles (summary from 2 replicates, each pooling 2 different mice) using the *Mreg* promoter as a bait in liver and kidney at ZT20. Data are shown in a window of +/− 250 kb from the bait (top). Profiles of differential contacts between liver and kidney (bottom) represented as signed log p-values (regularized t-test, positive values denote liver-enriched 4C contacts). Tracks of differential 4C contacts (signed log p-values), log2 fold change of DNase-I hypersensitivity between liver and kidney, and ChIP-exo of REV-ERBa and FOXA2. Regions of significant differential 4C contacts correspond to liver-specific DNase-I hypersensitive regions and REV-ERBa binding sites.

### Precise promoter-enhancer contacts underlie liver-specific mRNA rhythms

To test whether distinct chromatin loops would form at alternative nearby gene promoters with distinct temporal mRNA profiles, we searched for candidate genes where one promoter was rhythmically transcribed while the alternative one was nonrhythmic (Supplemental Figure S8). *Slc45a3* has two alternative transcripts using promoters 8 kb apart, with the shorter oscillating in the liver (rhythmic promoter, *Slc45a3*-short), while the longer not (flat promoter, *Slc45a3*-long). In kidney, neither *Slc45a3*-short nor *Slc45a3*-long 11showed robust transcript rhythms (Supplemental Figure S9). Targeting the *Slc45a3*-short promoter with 4C-Seq in liver and kidney showed liver-specific chromatin loops at three distal regions (two upstream, one downstream) (Figure 6A). Remarkably, these same regions did not form liver-specific chromatin loops with the *Slc45a3*-long promoter (Figure 6B), suggesting that promoters 8 kb apart can contact distinct enhancers. Overlaying 4C-Seq with DNase-Seq, we found that these chromatin loops link liver-specific DHSs specifically to the *Slc45a3*-short promoter (Figure 6C). These liver-specific DHSs are bound by liver-specific TFs, FOXA2 and ONECUT1, and clock TF, REV-ERBa, as shown in ChIP-Seq. Taken together, the 4C experiments suggest that enhancers can contact a rhythmic promoter while looping out nearby nonrhythmic alternative promoters, confining rhythmic enhancer activity to specific promoters (Figure 6D). Furthermore, rhythmically active enhancers can contact promoters in a tissue-specific manner. Thus, chromatin folding not only regulates tissue-specific rhythms, but also differentiates between closely spaced promoters to control rhythmic transcription with spatial precision.

**Figure 6.**
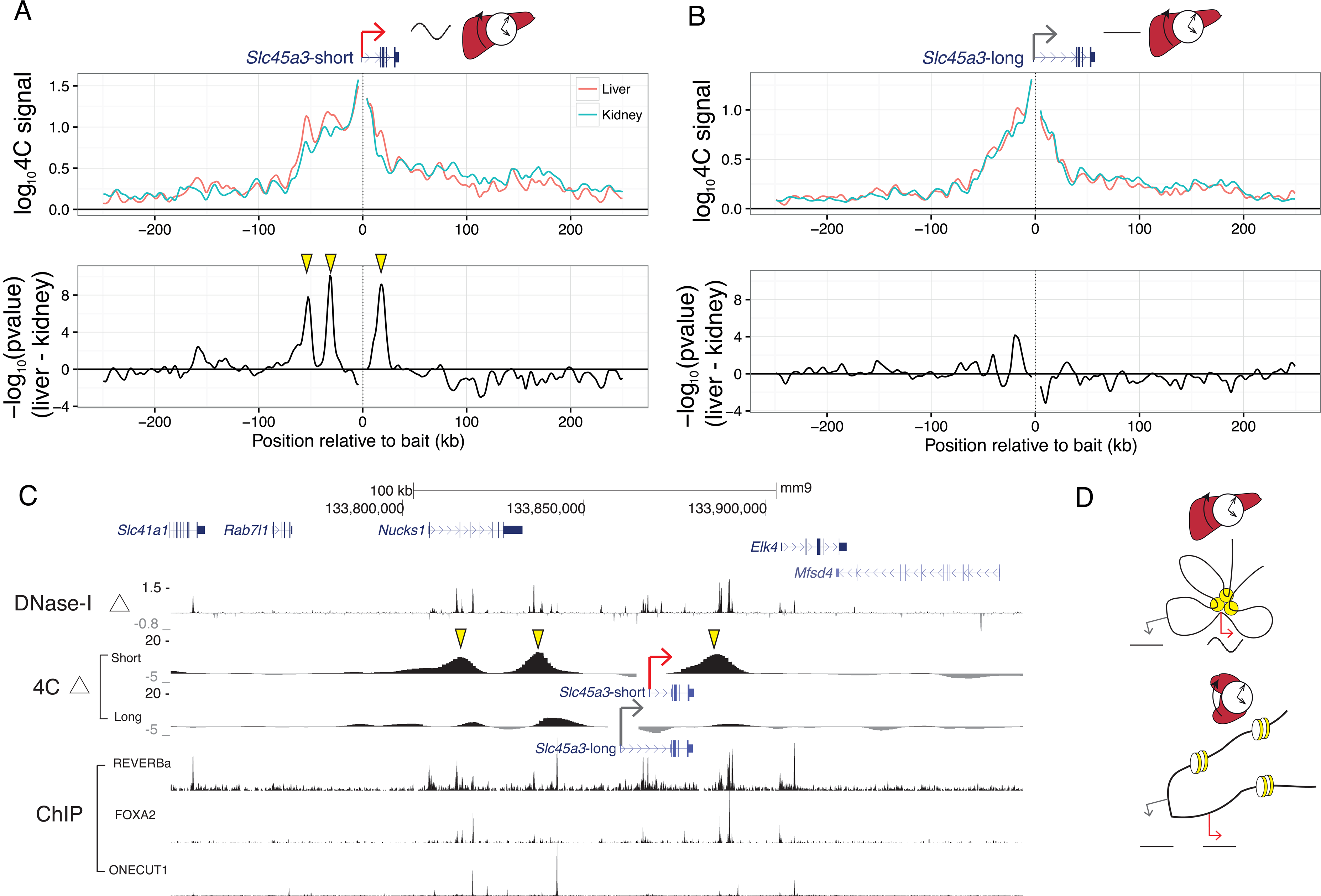
Precise promoter-enhancer contacts underlie liver-specific mRNA rhythms. (A,B) 4C-Seq profiles for the (A) *Slc45a3*-short and (B) *Slc45a3*-long isoforms within +/− 250 kb around baits targeting the two TSSs (top). Signed log p-values for differential contacts between liver and kidney (bottom) as in Figure 5B. TSSs for *Slc45a3*-short and *Slc45a3*-long are 8 kb apart. Yellow arrows denote liver-specific distal contacts found at the *Slc45a3*-short but absent at the *Slc45a3*-long TSS. (C) Differential 4C contacts (signed log p-values), log2 fold change of DNase-I hypersensitivity between liver and kidney, and ChIP-exo signal of REV-ERBa, FOXA2, and ONECUT1. Regions of significant differential contacts in *Slc45a3*-short correspond to liver-specific DNase-I hypersensitive regions. (D) Schematic model illustrating enhancer-promoter interactions in liver and kidney that may generate liver-specific rhythms. Yellow circles illustrate liver-active enhancers contacting the rhythmic promoter (red arrow) but not the alternative nonrhythmic promoter (grey). In kidney, the enhancer is not accessible and both promoters are nonrhythmic.

## Discussion

The mammalian genome encodes transcriptional programs that allow the molecular clock to robustly oscillate across diverse tissue transcriptomes while maintaining flexibility to regulate distinct clock outputs in different combinations of tissues. Here we identified two regulatory modes underlying tissue-specific transcript rhythms: (1) regulatory sequences can recruit individual TFs bearing rhythmic activity; (2) coordinated binding of clock and tissue-specific TFs can generate tissue-specific rhythms. Moreover, we found that clock and tissue-specific TFs bound at distal enhancers can be recruited to promoters through remarkably precise chromatin loops.

Several of our predictions of transcription regulators and regulated genes (e.g. EGR1, *Por, Upp2*) corroborated with previous analyses of independent datasets (Yan et al. 2008; Bozek et al. 2009; Bhargava et al. 2015). Further analysis incorporating outputs of enhancer activity, such as eRNAs (Fang et al. 2014), across multiple tissues may uncover additional rhythmically active regulators.

Co-localized binding of clock and tissue-specific TFs at enhancers provides a putative mechanism for the clock to regulate clock output genes in a tissue-specific manner. In mouse liver, clock TFs can co-localize with multiple liver-specific TFs, such as FOXA2 and ONECUT1, consistent with multiple liver TFs associating with liver-specific DHSs (Iwafuchi-Doi et al. 2016). Our findings are currently based on sequence-specific DNA binding of TFs, comparison of tissues, and ChIP-Seq datasets. Further mechanistic basis for the functional significance of co-localization could be gained, for example by using inducible knockout models for tissue-specific regulators. Moreover, the observed colocalization do not exclude other cooperative modes, such as tethering of REV-ERBa to ONECUT1 through protein-protein interactions (Zhang et al. 2015).

Our 4C analysis showed that chromatin looping might mediate interaction between clock and tissue-specific transcriptional programs, by recruiting clock-bound distal elements to promoters in a tissue-specific manner. Remarkably, such loops can surgically discriminate between nearby promoters as close as 8 kb apart, suggesting a way to separate the temporal regulation of neighboring promoters. A previous study applying 4C techniques to probe the contact landscape of a core clock gene enhancer proposed that cohesion-mediated promoter-enhancer looping can compartmentalize rhythmic gene expression within genomic regions spanning 150 kb (Xu et al. 2016). Here, chromatin interactions that differed between tissues were localized to a relatively small genomic region (<10 kb) proximal to the promoters (<100 kb). Future studies integrating temporal data across tissues with large-scale promoter-enhancer networks may reveal regulatory sequences that encode promoter-enhancer compatibility and elucidate whether this compatibility is tissue-specific (Li & Noll 1994; Merli et al. 1996; Zabidi et al. 2014; Nguyen et al. 2016).

Overall, this work proposed a role for newly identified rhythmic transcription factors and tissue-specific chromatin interactions in regulating tissue-specific rhythmic gene expression. While our work focused on transcriptional mechanisms, studying others mechanisms such as posttranscriptional, translational, and posttranslational processes using PRO-Seq, Ribo-Seq, and proteomics data may provide additional insights. Expanding our 24-hour analysis to 12-hour or other harmonics would broaden the view of tissue-specific temporal gene expression, but may require experimental designs of higher temporal resolution (Hughes et al. 2009; Krishnaiah et al. 2017). Tissues regulate dynamic physiological processes such as glucose homeostasis, lipid metabolism, and sodium homeostasis at different times of day. Thus, integrating the temporal axis into tissue-specific gene regulation offers an integrated understanding of how tissue physiology resonates with daily cycles in the environment.

## Materials and Methods

### Animal experiments

8-14 weeks old C57Bl/6 mice have been purchased from Charles River Laboratory. *Bmal1* KO mice have been previously described (Jouffe et al. 2013). Without further indications, mice are kept under 12 hours light/12 hours dark regimen and *ad libitum* feeding. All animal care and handling was performed according to the Canton de Vaud (Fred Gachon, authorization no VD 2720) laws for animal protection.

### RNA-Seq experiments and analysis

#### Processing

To complement the mouse liver WT and *Bmal1* KO RNA-Seq data (GSE73554) (Atger et al. 2015), transcriptomes of kidneys from *Bmal1* KO and WT littermates (12 hours light/12 hours regimen; night-restricted feeding) were measured following the same protocol as in (Atger et al. 2015). mRNA levels were quantified using kallisto version 0.42.4 (mm10) (Bray et al. 2015).

#### Global Temporal Variance

For each tissue, we estimated the contribution of temporal variance for each gene, broken down by its Fourier components. We calculated the background level assuming temporally unstructured data (white noise), whose magnitude (strength of the white noise) was estimated from the mean of squared magnitudes of Fourier coefficients that were not submultiples of 24 hours (i.e., the mean of 48, 16, 9.6, 6.9, 5.3, 4.4 hour components).

#### Model Selection

We fitted harmonic regression models that integrated temporal gene expression across different combinations of rhythms in different conditions (Atger et al. 2015). One difference from previous methods was that for comparing different models, we used a g-prior for the rhythmic parameters 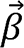 rather than BIC (Liang et al. 2008),

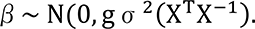

The hyperparameter g controls the spread of the prior over the models; as g increases, simpler models (such as tissue-wide model) are favored over more complex models (such as model with many divergent rhythms). We set g=1000, which we found to maximize temporal variations captured in the shared rhythms model while minimizing temporal variations captured in the flat model. The number of rhythmic combinations k scales as a function of the number of conditions n as *k*(*n*) = *B*_*n*+1_ where B is the Bell number used in combinatorial mathematics. See supplemental methods for details.

#### Complex singular value decomposition (SVD) representation of gene and tissue module

Gene expression over time and across tissues can be represented as a 3-dimensional array. However, since SVD of a tensor does not have all the properties of a matrix SVD, we first transformed the time domain to the frequency domain corresponding to 24-hour rhythms for all genes *g* and conditions *c:*

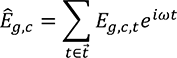

where *Ê*_*g,c*_ is a complex value representing the amplitude and phase of expression for gene *g* in condition *c* and *ω* = 2*π*/24.

The resulting matrix was decomposed using SVD and the first left -and right-singular values were visualized in separate polar plots. To ensure the first component recovered most of the original signal, the SVD representation was performed separately for each gene module identified by model selection.

### Predicting Activities of Transcriptional Regulators

#### Predictions of transcription factor binding site (TFBS)

For TFBS predictions near promoters, we used motevo version 1.03 (Arnold et al. 2012) to scan +/− 500 bp around the promoter. We used promoters (Balwierz et al. 2009) and weight matrices of transcription factors defined by SwissRegulon (Pachkov et al. 2013) (http://swissregulon.unibas.ch/fcgi/sr/downloads). For distal regions, we scanned the genome for TFBSs in 500 bp windows in genomic regions within 40 kb of an annotated gene.

#### Penalized regression model

We applied a penalized regression model as previously described (Balwierz et al. 2014) using an L2 penalty for penalization, which allows a direct estimate of the standard deviation. Rhythmic activities of transcription factor motifs were summarized using complex-valued singular value decomposition. We projected the activities to an amplitude and phase and calculated the zscore of the amplitude. We considered activities with zscore > 1.25 as rhythmic TF activities. Time of peak temporal activities of transcription factors were subtracted by 3 hours, to account for an average 3 hour shift between peak transcription and peak mRNA accumulation (Le Martelot et al. 2012).

#### Enrichment of pairs of motifs

We applied log-linear models to test for statistical significance between pairs of motifs across rhythmic versus nonrhythmic modules. For each motif, we ordered DHS sites by the posterior sitecount of the motif (decreasing order) and considered the motif to be present in the DHS site if the sitecount was in the top 300 (Myšičková et al. 2012). We considered liver-specific DHS sites that were annotated to a clock-dependent liver-rhythmic gene or to a nonrhythmic gene. For each annotated label and for each pair of motifs, we constructed a 2 by 2 contingency table by counting the number of DHS sites that contain one of the motifs, both motifs, or none, resulting in a 3-way contingency table (motif 1, motif 2, and annotated label). We assessed whether the resulting contingency table was statistically significant to a null model, where the null model was the expected counts if the pair of motifs were jointly independent on the rhythmicity.

### Chromatin conformation experiments and analysis

C57Bl/6 mice were sacrificed at ZT08 and ZT20 to extract liver and kidneys. Liver and kidney nuclei were prepared as previously described (Ripperger & Schibler 2006) with some minor changes. 4C-Seq assays were performed as in (Gheldof et al. 2012). See supplemental methods for details.

Raw read counts for each sample were normalized by library size by the sum of the read counts on the cis-chromosome (excluding 10 fragments around the bait). Read counts were log-transformed using the formula:

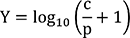

where p=500, the pseudocount.

The weighted linear model was fit locally around each fragment f. A Gaussian window centered on f was used to incorporate signal from neighboring fragments. The 4C-Seq fragment counts was modeled by the fragment effect i and tissue effect j.

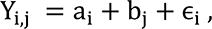

Where the weights of the linear model is defined as:

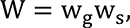

Where:
w_g_ is a Gaussian smoothing kernel (width σ _G_ = 2500 bp, centered on fragment f). w_s_ is the sample weight based on the number of non-zero values counts on fragment i, specifically, we used w_s_ = (0.5,1.5,2.5) for fragments with (0,1,2) finite counts out of the two replicates.

Differential contacts were estimated using t-statistics:

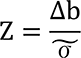

Where 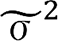 stands for the regularized sample variance:

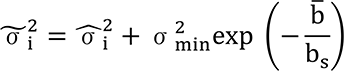

Where:

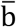 = the estimated signal across samples
b_s_ = log_10_(2)

## Data Access

Raw and processed data generated in this study are available in the Gene Expression Omnibus (GEO) database under accession number GSE100457.

## Acknowledgements

We thank Eric Paquet for critical reading and Saeed Omidi for help and discussions in bioinformatics. This work was supported by Swiss National Science Foundation Grant 31003A-153340, European Research Council Grant ERC-2010-StG-260667, and the Ecole Polytechnique de Lausanne. J.Y. benefits from the Natural Sciences and Engineering Research Council of Canada Postgraduate Studies Doctoral scholarship.

## Author Contributions

Conceptualization, J.Y., J.M., and F.N; Formal analysis, J.Y. and F.N.; Investigation, J.M., C.J., J.M., A.C.; Writing – Original Draft, J.Y., J.M., and F.N.; Writing - Review & Editing, J.Y., J.M., F.G., F.N.; Supervision, F.N. and F.G.; Funding Acquisition, F.N. and F.G.

